# The network underlying human higher-order motor control: Insights from machine learning-based lesion-behaviour mapping in apraxia

**DOI:** 10.1101/536391

**Authors:** Christoph Sperber, Daniel Wiesen, Georg Goldenberg, Hans-Otto Karnath

## Abstract

Neurological patients with apraxia of pantomime provide us with a unique opportunity to study the neural correlates of higher-order motor function. Previous studies using lesion-behaviour mapping methods led to inconsistent anatomical results, reporting various lesion locations to induce this symptom. We hypothesised that the inconsistencies might arise from limitations of mass-univariate lesion-behaviour mapping approaches if our ability to pantomime the use of objects is organised in a brain network. Thus, we investigated apraxia of pantomime by using multivariate lesion behaviour mapping based both on support vector regression and sparse canonical correlations in a sample of 130 left-hemisphere stroke patients. Both multivariate methods identified multiple areas to underlie high-order motor control, including inferior parietal lobule, precentral gyrus, posterior parts of middle temporal cortex, and insula. Further, long association fibres were affected, such as the superior longitudinal fascicle, inferior occipito-frontal fascicle, uncinated fascicle, and superior occipito-frontal fascicle. The findings thus not only underline the benefits of multivariate lesion-behaviour mapping in brain networks, but they also uncovered that higher-order motor control indeed is based on a common anatomical network.

## Introduction

Following brain damage primarily of the left hemisphere, patients can suffer from high-order motor disorders not caused by primary motor or sensory deficits (Heilman & Rothi, 1993). These disorders are summarised under the umbrella term ‘apraxia’ (Wheaton & Hallett, 2007; Goldenberg, 2011) and can include, for example, our ability to imitate or execute gestures (De Renzi et al., 1980; Goldenberg, 1996; Rumiati et al., 2009; Mengotti et al., 2015), to perform motor imagery (Ochipa et al., 1997; Buxbaum et al., 2005), or to mechanically reason (Goldenberg & Hagmann, 1998; Baumard et al., 2014). A prominent disorder in this field is apraxia of pantomime (hereinafter simply referred to as ‘apraxia’) in which patients fail to pantomime the use of a common tool as if they hold the tool in hand, while they are typically able to use the real tool with less or no errors (De Renzi et al., 1982; Goldenberg & Hagmann, 1998; Wada et al., 1999; Lausberg et al., 2003; Goldenberg et al., 2004; Laimgruber et al., 2005; Sperber et al., 2018).

Anatomo-behavioural studies with lesion-behaviour mapping offer a unique opportunity to study the brain architecture underlying these higher order motor skills. Multiple such studies investigated the neural correlates of apraxia (for a review see Niessen et al., 2014). Their results, however, were inconsistent. Most frequently, apraxia was associated with lesions to the inferior parietal lobe or adjacent parietal regions (Halsband et al., 2001; Buxbaum et al., 2003, 2005; Weiss et al., 2008; Hoeren et al., 2014; Goldenberg & Randerath, 2015). On the other hand, several studies found lesions in the inferior frontal gyrus to induce apraxia (Goldenberg et al., 2007; Manuel et al., 2013; Weiss et al., 2016). Besides, regions such as the insula (Goldenberg et al., 2007; Hermsdörfer et al., 2013; Hoeren et al., 2014), premotor and precentral areas (Weiss et al., 2016), and the middle temporal gyrus (Manuel et al., 2013) were also reported to be critical. Interestingly, results in most studies were limited to only a single or few of these areas.

There are many possible methodological reasons for these inconsistencies (for a review see Sperber & Karnath, 2018), including, e.g., time since stroke and differences in apraxia assessment and rating. Another potential major source for heterogeneous results could be the general analysis approach of the above mentioned studies. While they used different analysis techniques – such as voxel-based lesion behaviour mapping (VLBM; Manuel et al., 2013; Hoeren et al., 2014; Goldenberg & Randerath, 2015; Weiss et al., 2016; Finkel et al., 2018), subtraction plots (Goldenberg et al., 2007; Weiss et al., 2008; Hermsdörfer et al., 2013), or region-of-interest analyses (Halsband et al., 2001) – all studies followed a univariate approach. In such univariate approach for topographical analyses, each voxel in an image is analysed independently of other voxels. Univariate methods such as mass-univariate VLBM, however, can fail to identify neural correlates of pathological behaviour if behaviour is organised in larger modules or networks (Rorden et al., 2009; Mah et al., 2014; Zhang et al., 2014; Gajardo-Vidal et al., 2018; Pustina et al., 2018; Sperber et al., in press). The above mentioned anatomo-behavioural studies on apraxia thus might have been unable to gain full insight to the neural correlates of human higher-order motor control, and instead only identified single components of a possible network.

Previous anatomical findings point at the existence of a brain network to underlie apraxia. First, the above reported findings from mass-univariate studies already identified different areas that are spread across the brain, and that might be cortical nodes of such network. This point is also supported by componential analyses, that investigated multiple different sub-symptoms of pantomime (Manuel et al., 2013; Finkel et al., 2018). These studies found different areas to underlie behavioural components of apraxia, which could be part of a network for motor control. Second, a role of white matter fibre connections between these areas has been found not only in some of the above mentioned studies (e.g. Goldenberg et al., 2007; Manuel et al., 2013; Hoeren et al., 2014), but also in single cases (Kertesz and Ferro, 1984) and by a fMRI-DTI study (Vry et al., 2015). Thus, damage to a brain network of multiple cortical regions that are connected by white matter fibres likely underlies apraxia (see also Niessen et al., 2014; Vry et al., 2015; Finkel et al., 2018; Buxbaum & Randerath, 2018), which already has been postulated in classical models of praxis skills by Liepmann and Geschwind (see Buxbaum & Randerath, 2018).

In recent years, multivariate lesion-behaviour mapping methods based on machine learning have been developed (Smith et al., 2013; Mah et al., 2014; Zhang et al., 2014; Yourganov et al., 2015; Pustina et al., 2018). Multivariate lesion-behaviour mapping includes multiple variables – e.g., the lesion status of multiple voxels or regions of interest – into one single model. Simulation studies have shown that such methods perform better than traditional VLBM tools if damage to multiple brain regions can induce a particular symptom (Mah et al., 2014; Zhang et al., 2014; Pustina et al., 2018). Thus, if indeed a network underlies neuropsychological deficits in the ability to pantomime, multivariate analyses might resolve the inconsistencies in the field of apraxia. To test this hypothesis, we reanalysed a large sample of left hemisphere stroke patients using both univariate and multivariate lesion behaviour mapping methods. As multivariate methods, we chose two different approaches, one based on support vector regression, and another based on sparse canonical correlations.

## Methods

### Subjects

We retrospectively analysed data of 130 left brain damaged patients (mean age = 56.5 ± 12.3 years; range 26-83) that had been admitted to the Neuropsychological Department of the Bogenhausen Hospital in Munich. Demographic and clinical data are given in Table 1. The patients have been investigated in two previous studies (Goldenberg et al., 2007; Goldenberg & Randerath, 2015). All patients had a first ever left hemisphere stroke at least three weeks before the examination. Neuropsychological examination and imaging were part of clinical protocols at the Bogenhausen Hospital. Patients consented to the scientific re-use of their data; the study has been performed in accordance with the ethical standards laid down in the revised Declaration of Helsinki.

**Table 1:** Demographic and clinical data. Demographic and clinical data of all 130 patients. Pantomime was tested using a 20-item test (Goldenberg et al., 2003, 2007); imitation of hand and finger gestures was assessed using a 10-item test each (Goldenberg, 1996). Patients were considered to have defective pantomime when they scored below a cutoff of 45 points. This cutoff was defined with a sample of 49 healthy control subjects (Goldenberg et al., 2007). Maximum score obtainable in the pantomime task was 55 points (cutoff < 45) and 20 points in the imitation tasks (finger posture imitation cutoff < 17; hand posture imitation cutoff < 18). Aphasia was classified according to the Aachen Aphasia Test. Numbers in parentheses indicate standard deviations.

|  | Pantomime normal | Pantomime defective |
| --- | --- | --- |
| Age (years) | 55.4 (11.4) | 57.7 (13.1) |
| Ischemia/haemorrhage | 54/14 | 49/13 |
| Lesion size (mm<sup>3</sup>) | 79.2 (58.4) | 111.1 (71.2) |
| Time since lesion (weeks) | 14.2 (18.6) | 14.9 (17.9) |
| Aphasia classification | 6 none/ 12 global/ 9<br>broca/ 16 wernicke/ 10<br>amnesic/ 15 other | 0 none/ 27 global/ 4<br>broca/ 12 wernicke/ 8<br>amnesic/ 11 other |
| Hemiparesis present (%) | 44.1 | 61.3 |
| Imitation finger (test score) | 17.8 (2.9) | 14.6 (4.4) |
| Imitation finger defective (%) | 32.4 | 59.7 |
| Imitation hand (test score) | 16.8 (3.8) | 14.4 (4.6) |
| Imitation hand defective (%) | 44.9 | 67.7 |
| Pantomime (test score) | 49.2 (3.0) | 29.6 (10.6) |

### Neuropsychological examination

Pantomime of tool use was assessed with a 20-item test (Goldenberg et al., 2003, 2007). Patients were asked to pantomime the use of common tools as if they hold the actual tool in hand, but without the object being actually present. For each item the examiner named an action and the corresponding tool and simultaneously showed a picture of the object (e.g., ‘How do you brush your teeth with a tooth brush?’ while showing a picture of a tooth brush). The picture was removed before the patient initiated the task. Patients with hemiparesis were instructed to use the left, ipsilesional hand to perform the task. To ensure that patients did understand the task, practice items were performed and patients that did not understand the task (e.g., when a patient drew the outlines of the tool on the table instead of performing pantomime) were excluded. For each of the 20 items one point was scored for the correct grip and finger posture and a maximum of one to three points for aspects such as movement amplitude, trajectories, or hand position in relation to the own body. Maximum score was 55 points. The inter-rater reliability was previously found to be very satisfying both for the number of correct features per item (kappa = 0.61) and for the total test score (kappa = 0.94; Goldenberg et al., 2003). Furthermore, all patients were tested for aphasia with the Aachen Aphasia Test (Huber et al., 1983) and for apractic deficits in imitation of postures with the fingers or the hand (Goldenberg, 1996).

### Imaging and lesion mapping

Structural imaging was acquired either by MRI (n = 118) or CT (n = 12) on average 14.6 weeks (SD 18.2) after stroke-onset. The interval between imaging and neuropsychological examination was maximally three weeks. Lesions were manually mapped on transversal slices of the T1-weighted ‘ch2’ template scan from the Montreal Neurological Institute using MRIcro software (Rorden & Brett, 2000; http://people.cas.sc.edu/rorden/mricro/index.html). The ‘ch2’ template is oriented to fit Talairach space (Talairach & Tournoux, 1998) and is distributed with the MRIcro software. Lesions were mapped on a fixed set of twelve slices with z-coordinates −40, −32, −24, −16, −8, 0, 8, 16, 24, 32, 40, and 50 by using the closest matching or identical transversal slice found in the imaging. Modern normalisation techniques were not applicable here, because the original scans were not available; see the discussion section for further details. A topography of all lesions is shown in Figure 1A. To obtain an estimate for lesion size for the demographic data (Table 1), lesions were interpolated by converting each individual slice into a volume of 8mm thickness.

**Figure 1:**
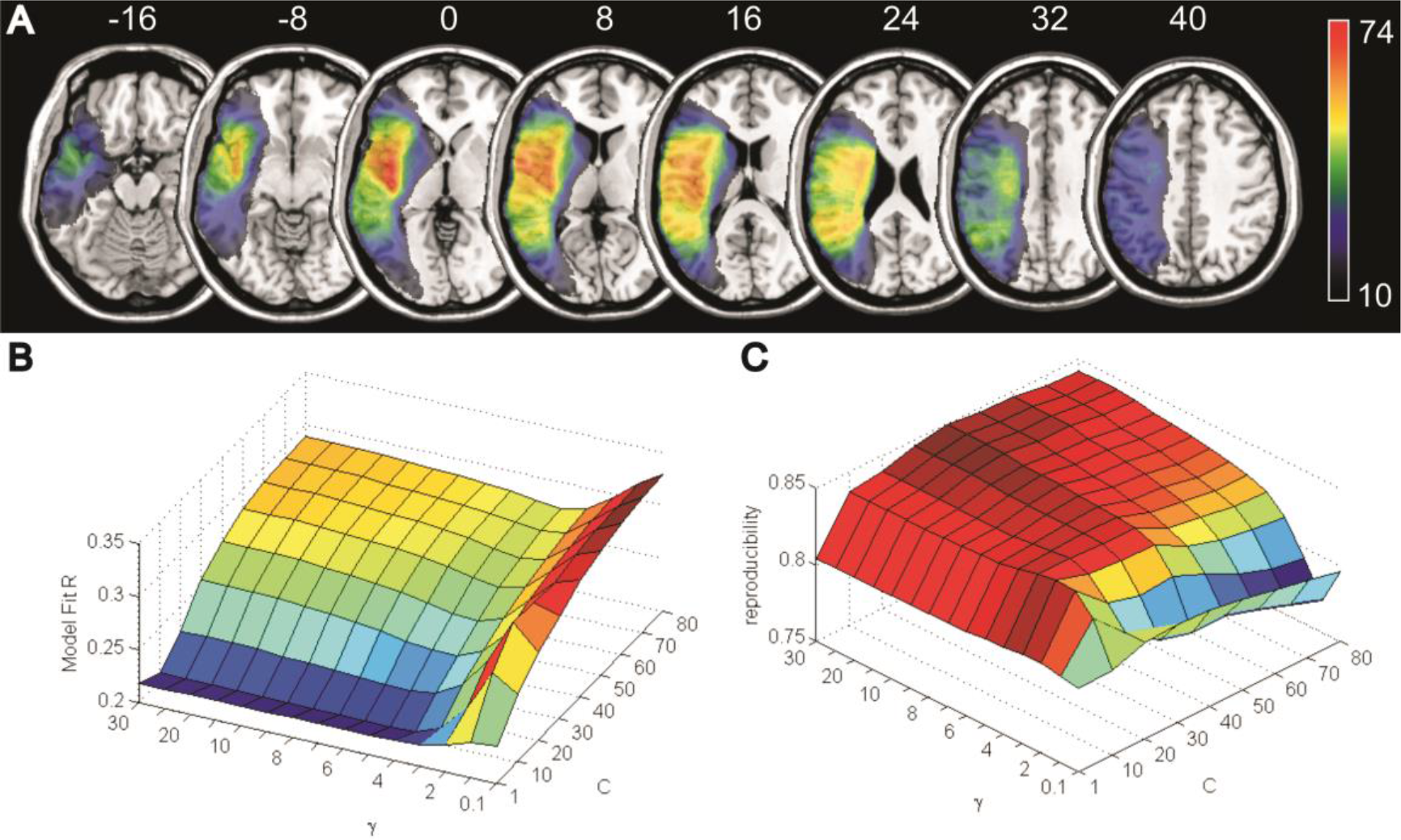
Topography of brain lesions and hyper-parameter optimisation for the SVR-LSM. **(A)** Lesion topography of all 130 brain lesions with colour-coding that depicts the number of overlaying lesions per voxel. Only voxels affected in at least ten patients, i.e. voxels included in the multivariate analysis, are shown. Numbers above the slices indicate z-coordinates in MNI space. **(B)** Results of the hyper-parameter optimisation by grid search for C and γ in regards to model Fit R and **(C)** reproducibility (see Rasmussen et al., 2012; Zhang et al., 2014) are plotted for the a priori set of chosen parameters.

### Univariate lesion behaviour mapping

As a baseline for comparison with the multivariate lesion behaviour mapping methods we conducted several analyses using mass-univariate VLBM. First, we conducted VLBM analyses on the continuous apraxia scores using t-tests with correction for multiple comparisons both by false discovery rate (FDR) thresholding and permutation-based thresholding with 4000 permutations. We chose to include both methods, as FDR correction is often used, but permutation thresholding can be considered to be a better approach in VLBM (Karnath et al., 2018; Pustina et al., 2018). Both analyses were once conducted with control for lesion size and once without. All analyses were performed at the p < 0.05 level using NiiStat software (https://github.com/neurolabusc/NiiStat), lesion size was controlled for by the nuisance regression approach implemented in NiiStat. Only voxels affected in at least 10 patients were included into the analyses.

### Multivariate lesion behaviour mapping by SVR-LSM

Multivariate lesion behaviour mapping (MLBM) was first performed using support vector regression (SVR; Vapnik, 1995; Drucker et al., 1996), which is a multivariate regression method based on machine learning. This method is an extension of support vector machines (Cortes and Vapnik, 1995). SVR is able to model continuous variables and has been successfully implemented in SVR-based lesion symptom mapping (SVR-LSM) to map lesion-behaviour relationships with high resolution on a whole-brain voxel-level (Zhang et al., 2014; Mirman et al., 2015b; Fama et al., 2017; Griffis et al., 2017; Ghaleh et al., 2018; DeMarco & Turkeltaub, 2018; Chen et al., 2018; Sperber et al., in press). Most importantly, SVR-LSM was validated in a set of simulation studies for simple brain networks (Zhang et al., 2014). Thus, at least in such artificial situations, voxel-wise SVR-LSM was empirically proven to be able to identify critical brain regions assembled in networks.

The analysis was performed with MATLAB 2016a and libSVM (Chang and Lin, 2011). We modified a publicly available collection of scripts (https://github.com/yongsheng-zhang/SVR-LSM) used in the study by Zhang et al. (2014). Among others, we adopted algorithms for control for lesion size and for the derivation of a topography from SVR parameters. Central functions of the toolbox were not altered, and a control analysis with the original scripts replicated the results. The detailed methodological process and theoretical background is described in Zhang et al. (2014). In short, binary lesion images were vectorised and then normalised to have a unit norm. This procedure, called direct total lesion volume control, provides a control for the effect of lesion size. Lesion data and behavioural data were then further processed to fit the required data format of the libSVM toolbox. Using default libSVM options, an epsilon-SVR with radial basis function kernel was performed. The β-parameters obtained from the SVR were then remapped onto a three-dimensional brain topography. To assess statistical significance, a permutation approach has been chosen. By randomisation of behavioural scores, a large number of SVR-β-maps were generated and voxel-wise significance of β-parameters could be derived. The topography was computed using permutation testing with 4000 permutations, and only voxels with at least ten lesions were tested. To control for multiple comparisons, which is required in SVR-LSM (Sperber et al., in press), we applied false discovery rate correction (FDR; Benjamini and Yekutieli, 2001) at q = 0.1. Note that correction by FDR has some limitations in SVR-LSM; however, there is currently no consensus on clearly superior alternatives (for further discussion see Sperber et al., in press; but also de Marco & Turkeltaub, 2018). To obtain an optimal model, we performed an optimisation for hyper-parameters C and γ via grid search. The range of investigated parameters was chosen as in the study by Zhang et al. (2014): C = 1, 10, 20, 30, 40, 50, 60, 70, 80, and γ = 0.1, 1, 2, 3, 4, 5, 6, 7, 8, 9, 10, 15, 20, 25, 30. Using a five-fold cross-validation as in the study by Zhang et al. (2014), we evaluated both model fit and reproducibility of each parameter set (see Rasmussen et al., 2012; Zhang et al., 2014). Five-fold cross-validation was performed by splitting the data in five equally sized sub-samples. Reproducibility was assessed by modelling each set of 4 sub-samples taken together and computing the average correlation between β-parameters of the resulting weight maps. Model fit was tested by training a model on each set of 4 sub-samples taken together, then predicting the last fifth of the data with the model. Average explained variance was then computed to obtain model fit. Resulting topographies were interpreted according to the AAL atlas (Tzourio-Mazoyer et al., 2002) for grey matter regions. Due to marked differences between histology-based and tractography-based white matter atlases (de Haan & Karnath, 2017), white matter regions were interpreted according to both a probabilistic cytoarchitectonic fibre tract atlas (Bürgel et al., 2006) and a tractography-based probabilistic atlas (Thiebaut de Schotten et al., 2011). For the white matter atlases, overlay of the thresholded statistical map with the probabilistic map at p ≥ 0.3 was identified.

### Multivariate lesion behaviour mapping by sparse canonical correlations

MLBM was also performed with a recently established method based on sparse canonical correlations (Avants et al., 2014; Pustina et al., 2018; Thye & Mirman, 2018). Like SVR-LSM, this method performs a voxel-wise lesion behaviour mapping with continuous behavioural scores. A major difference to SVR-LSM is that SCCAN does not interpret voxel-wise weights based on a permutation test. Instead, SCCAN is based on an optimization procedure that gradually builds a multivariate model based on anatomical data from multiple voxels, which maximizes correlations with behavioural scores. Individual features that contributed to the model are not assessed for statistical significance. Instead, the correlation coefficient in the final model obtained from this procedure is statistically tested by assessing the model’s predictive performance in a cross-validation procedure. As this is done for a single global model, no multiple comparisons are performed. Like SVR-LSM, SCCAN was validated in a simulation study that has shown that the approach is indeed superior to univariate methods in detecting neural correlates consisting of multiple regions (Pustina et al., 2018).

The analysis was performed with the LESYMAP toolbox v.0.0.9008 (https://github.com/dorianps/LESYMAP) in R software using ANTsR. A hyper-parameter in this approach, the so-called sparseness value, is automatically optimized by 4-fold cross-validation. Only voxels affected in at least 10 patients were included in the analysis.

## Results

### Univariate lesion behaviour mapping

Voxel-based lesion behaviour mapping using continuous apraxia scores without control for lesion size did not yield any significant results for FDR correction, and only ten significant voxels for permutation thresholding. These voxels were located in the posterior periventricular white matter. Utilising a control for lesion size, neither VLBM with FDR nor VLBM with permutation thresholding yielded significant results.

### Multivariate lesion behaviour mapping using SVR-LSM

Like in the original implementation of SVR-LSM (Zhang et al., 2014), the grid search showed that model fit and reproducibility generally were diametrical (Fig. 1B and C). No objective guidelines for hyper-parameter selection available are available, but the ideal model should provide a fair model fit while maximising reproducibility (Rasmussen et al., 2012). Importantly, model fit is not supposed to be maximised in SVR-LSM. First, the statistical map is derived from voxel-wise β-weights via permutation testing and thus stability of these weights has highest priority (Rasmussen et al., 2012). Second, behavioural variables do not exclusively depend on anatomical information, (see Price et al., 2017), and purely anatomical models might – for many behavioural variables - per se be deemed to not achieve a high model fit (see Sperber et al., in press for further discussion). This is not a problem in the present analysis, as SVR-LSM only intends to capture and map the variance that is actually present in anatomical data. We chose C = 30 and γ = 4, as this set of hyper-parameters provided both a comparatively satisfying model fit (r = .24) and high reproducibility (reproducibility = .84).

The SVR-LSM topography (Fig. 2) revealed several highly significant left parietal, temporal, frontal, and subcortical regions to underlie apraxia. With correction for multiple comparisons by FDR, however, no significant voxels remained. Yet, at an uncorrected p-level of p < 0.01, 3.8% of all tested voxels became significant, which is much higher than the 1.0% of significant voxels that would be expected in data without any signal (see also Sperber et al., in press). These findings hint at large amounts of low signal present in the data. Therefore, we here decided post-hoc to report uncorrected results at p < 0.01. Note that results thus should be interpreted with caution, and only in comparison with the other multivariate analysis. Table 2 lists affected grey and white matter structures. Large areas of significant voxels were found in the inferior parietal lobule, including angular and supramarginal gyri. In frontal areas, we found a large cluster in precentral gyrus and only a small cluster in the periventricular white matter. Furthermore, significant clusters were found in middle temporal and occipital gyri, amygdala, and hippocampus. Large significant clusters were also found in the white matter. With reference to the histology-based atlas, these included fibre tracts such as the superior longitudinal fascicle, inferior occipito-frontal fascicle, and superior occipito-frontal fascicle. With reference to the tractography-based atlas, they further included the inferior longitudinal fascicle.

**Figure 2:**
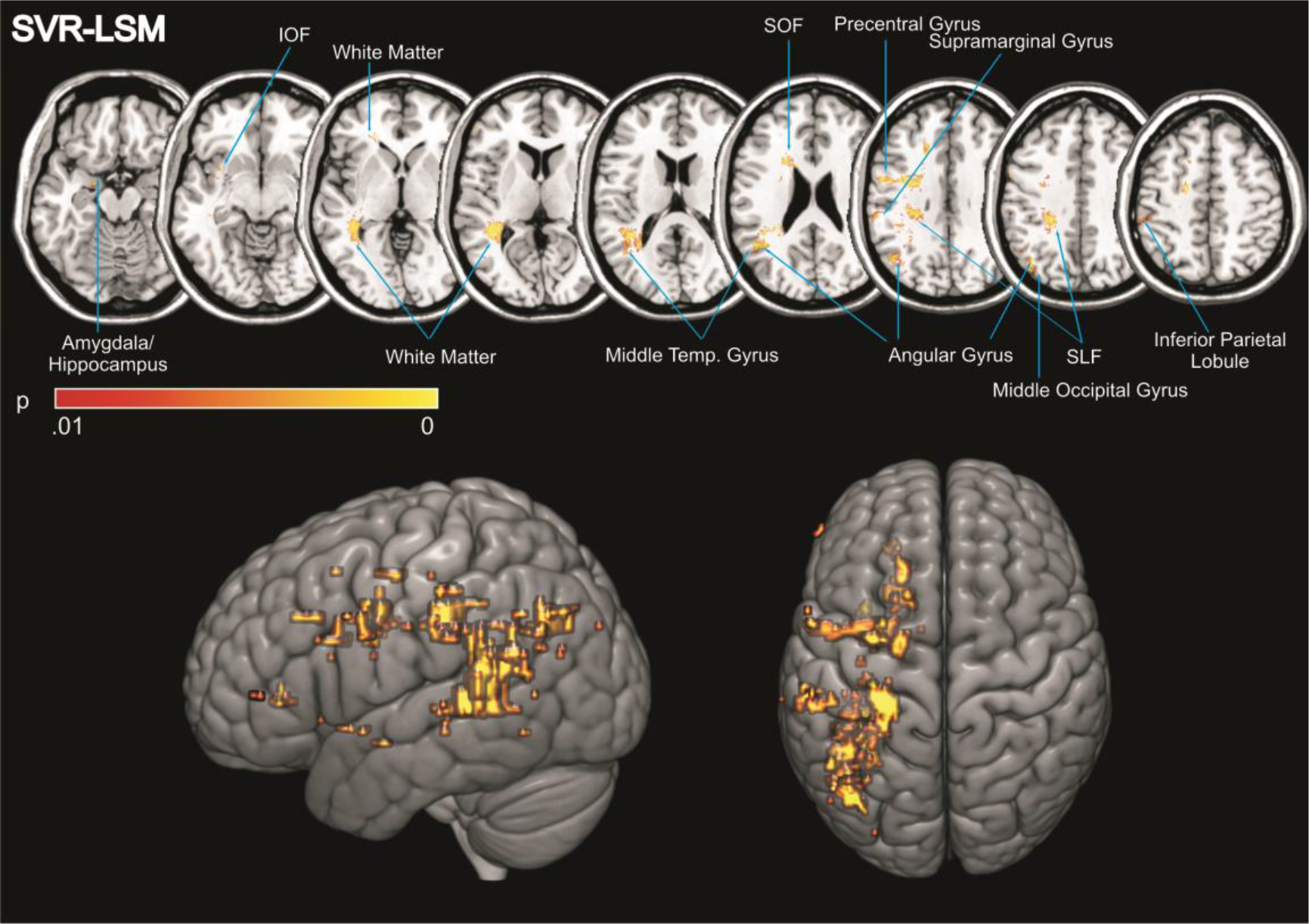
Results of multivariate lesion behaviour mapping by SVR-LSM. Permutation-thresholded β-map of voxel-wise SVR-LSM at p < 0.01 uncorrected, illustrating the anatomical regions significantly associated with apraxia of pantomime. Significant clusters were interpreted according to the AAL atlas (Tzourio-Mazoyer et al., 2002) for grey matter regions and to the probabilistic cytoarchitectonic fibre tract atlas (Bürgel et al., 2006) for white matter structures. Abbreviations: SLF – superior longitudinal fascicle; SOF – superior occipitofrontal fascicle; IOF – inferior occipitofrontal fascicle. Lower panels show full three-dimensional renderings of the same map using the 3D-interpolation algorithm provided by MRIcron (http://people.cas.sc.edu/rorden/mricron/index.html).

**Table 2:** Topological grey and white matter analysis for voxel-wise SVR-LSM. Topological analysis of grey and white matter structures covered by the voxel-wise SVR-LSM map thresholded at p = 0.01, uncorrected for multiple comparisons (see Fig. 2A). For grey matter structures, left hemispheric regions taken from the AAL Atlas (Tzourio-Mazoyer et al., 2002) with at least 1% affection are reported. For white matter structures, ROIs in a probabilistic histological atlas (Bürgel et al., 2006) and in a tractography-based atlas (Thiebaut de Schotten et al., 2011) were defined at a probability of p ≥ .3 to obtain binary maps. Of these binary maps, only left hemisphere parts were considered (MNI-coordinate x < 91). Further, for both grey and white matter atlas ROIs only z-slices that were part of the statistical analysis were considered.

| Grey matter structure | Percent affected | White matter structure (Jülich Histological Atlas) | Percent affected |
| --- | --- | --- | --- |
| Angular gyrus | 17.2 | Sup. longitudinal fasc. | 27.1 |
| Amygdala | 9.0 | Sup. occ.-frontal fasc. | 9.5 |
| Precentral gyrus | 5.2 | Corticospinal tract | 7.6 |
| Supramarginal gyrus | 4.2 | Optic radiation | 5.4 |
| Middle temporal gyrus | 1.6 | Callosal body | 4.7 |
| Middle occipital gyrus | 1.6 | Inf. occ.-frontal fasc. | 2.6 |
| Hippocampus | 1.4 |  |  |
| Inferior parietal lobe | 1.2 |  |  |
|  |  | White matter structure (Catani DTI Atlas) | Percent affected |
|  |  | Posterior segment (Arcuate) | 39.6 |
|  |  | Long segment (Arcuate) | 22.9 |
|  |  | Arcuate | 18.4 |
|  |  | Sup. longitudinal fasc.2 | 16.7 |
|  |  | Optic radiation | 14.4 |
|  |  | Inf. longitudinal fasciculus | 13.5 |
|  |  | Cortico ponto cerebellum | 7.0 |
|  |  | Internal capsule | 6.7 |
|  |  | Corticospinal tract | 6.6 |
|  |  | Inf. occ.-frontal fasc. | 5.3 |
|  |  | Anterior segment (Arcuate) | 2.9 |
|  |  | Corpus callosum | 2.9 |
|  |  | Uncinate | 1.7 |

### Multivariate lesion behaviour mapping using SCCAN

The MLBM using SCCAN identified multiple clusters that were included into the multivariate model (Fig. 3), and performed with a cross-validation correlation of r = 0.32 (p = 0.00018) and an optimal sparseness of 0.33. Table 3 lists affected grey and white matter structures. Note again that SCCAN does not rely on multiple statistical tests. Thus, correction for multiple comparisons is not required and present results on apraxia are significant.

**Figure 3:**
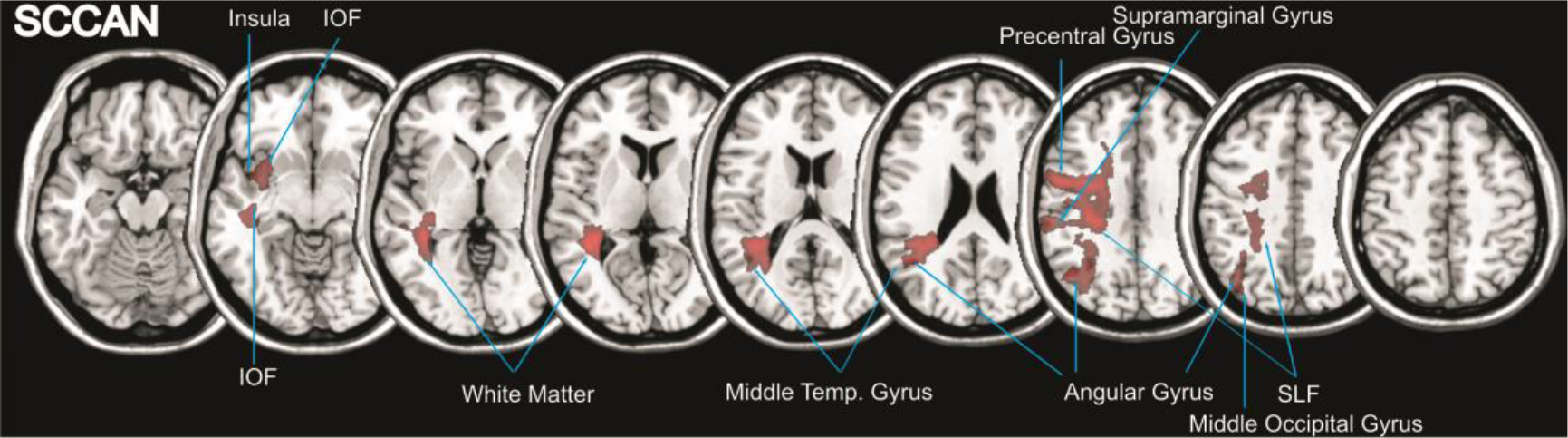
Results of voxel-wise multivariate lesion behaviour mapping by SCCAN. Topography of areas associated with apraxia in the SCCAN analysis. Clusters were interpreted with the same atlases as in Figure 2.

**Table 3:** Topological grey and white matter analysis for SCCAN. Topological analysis of grey and white matter structures covered by SCCAN map (see Fig. 3). Areas where interpreted following the same strategy as in table 2.

| Grey matter structure | Percent affected | White matter structure (Jülich Histological Atlas) | Percent affected |
| --- | --- | --- | --- |
| Angular gyrus | 30.4 | Inf. occ.-frontal fasc. | 52.4 |
| Supramarginal gyrus | 18.3 | Sup. occ.-frontal fasc. | 49.4 |
| Middle temporal pole | 17.8 | Uncinate fascicle | 27.0 |
| Precentral gyrus | 14.2 | Corticospinal tract | 25.2 |
| Middle occipital gyrus | 6.2 | Optic radiation | 13.0 |
| Insula | 5.4 | Callosal body | 11.1 |
| Superior temporal pole | 3.5 | Acoustic radiation | 10.2 |
| Middle temporal gyrus | 2.8 | Sup. occ.-frontal fasc. | 6.8 |
| Inferior parietal lobe | 2.3 |  |  |
| Hippocampus | 2.2 |  |  |
| Superior temporal gyrus | 2.1 |  |  |
| Postcentral gyrus | 1.7 |  |  |

| White matter structure (Catani DTI Atlas) | Percent affected |
| --- | --- |
| Posterior segment (Arcuate) | 65.4 |
| Sup. longitudinal fasc. 2 | 61.1 |
| Arcuate | 39.3 |
| Long segment (Arcuate) | 36.0 |
| Inf. longitudinal fasciculus | 28.5 |
| Optic radiation | 28.0 |
| Corticospinal tract | 17.7 |
| Inf. occ.-frontal fasc. | 16.1 |
| Internal capsule | 15.8 |
| Cortico-ponto-cerebellar | 14.8 |
| Uncinate | 10.7 |
| Anterior segment (Arcuate) | 8.7 |
| Corpus callosum | 6.5 |
| Fornix | 3.2 |

The SCCAN analysis also identified parietal, temporal, frontal, and subcortical regions to underlie apraxia. Largest clusters were found in the inferior parietal lobule, including the angular and supramarginal gyri. In the frontal lobe, again, the precentral gyrus was found. In the temporal lobe, significant regions included the middle and superior gyri and the temporal pole. This cluster reached into the middle occipital gyrus. Different to SVR-LSM, SCCAN also revealed the insula. In the subcortical white matter, SCCAN revealed qualitatively more significant areas than SVR-LSM. According to the histology-based atlas such regions included the superior longitudinal fascicle, inferior occipito-frontal fascicle, and superior occipito-frontal fascicle, but also the uncinate fascicle. According to the tractography-based atlas the topography further included the inferior longitudinal fascicle.

## Discussion

In a large sample of 130 left brain damaged patients, we investigated apraxia of pantomime by using multivariate lesion behaviour mapping. A main advantage of MLBM is that the role of different brain regions can be investigated in combination. Accordingly, we identified multiple cortical regions and white matter fibres underlying human higher-order motor control. Different parts of this network have been observed in previous studies in isolation. This has contributed to the inconsistent reports and conclusions on the neural representation of apraxia. Our multivariate analysis uncovered that these regions belong to a common network underlying higher-order motor control.

In contrast, mass-univariate analyses failed to find larger signal. This underlines the limitations of mass-univariate approaches in apraxia. The most likely reason for the inability of mass-univariate approaches to identify the apraxia network is the ‘partial injury problem’. This methodological problem is inherent to several lesion analysis methods including VLBM (Rorden et al., 2009; for review Karnath et al., 2018 or Sperber et al., in press). Imagine a simple brain network consisting of two distinct areas A and B. If damage to either area A or area B can induce the same symptom, patients showing the deficit may exist that have damage to area A, but not area B, and vice versa. Such patients will be used as counter examples in the voxel-wise statistical tests and statistical power to detect the neural correlates of the symptom is therefore reduced. Thus, mass-univariate methods can fail to identify a brain network in parts or – as in the present study – in whole. Importantly, previous mass-univariate studies thus did not provide ‘wrong’ results, but simply were unable to identify all critical brain regions involved at once. A plethora of factors might play a role in what regions of a network are found in a univariate analysis. Among those are several factors regarding patient selection, patient assessment, and study design (see Sperber & Karnath, 2018). But even taking these factors aside, significant results can be driven by sub sets of patients (Gajardo-Vidal et al., 2018), which might be especially a problem in smaller studies with lower power (Lorca-Puls et al., 2018). This also explains why the present study provided different results than two previous studies that partially used the same data (Goldenberg et al., 2007; Goldenberg & Randerath, 2015).

Another reason for the failure of mass-univariate approaches in apraxia in the present study could be low signal in the data. An argument in favour of this assumption is the observation that even if signal from all over the brain was modelled at once by SVR-LSM, the results did not survive correction for multiple comparisons. However, the second MLBM analysis by SCCAN – which does not rely on multiple statistical comparisons – found topographical results that were highly similar to the results of uncorrected SVR-LSM. Both analyses associated apraxia with the angular and supramarginal gyri, middle temporal gyrus, and precentral gyrus. Furthermore, both analyses found large clusters in white matter, which included fibre tracts like superior longitudinal fascicle, inferior occipito-frontal fascicle, and superior occipito-frontal fascicle. While the correspondence of both topographies was high, there were still small discrepancies. Notably, SCCAN found a qualitatively more pronounced role of white matter tracts, and also signal in the insula, while the uncorrected SVR-LSM found some small clusters in periventricular frontal white matter. Interestingly, involvement of the insula in apraxia has been reported by several previous studies (Goldenberg et al., 2007; Hermsdörfer et al., 2013; Hoeren et al., 2014).

A major advantage of lesion studies in comparison with functional studies based e.g. in fMRI is that the role of fibre tracks can be investigated. Both MLBM analyses of the present study found large clusters in white matter tracts. These included the superior longitudinal fascicle, the inferior occipito-frontal fascicle, the uncincate fascicle, and the superior occipito-frontal fascicle. These white matter fibres constitute a perisylvan network that connects frontal, temporal, and parietal brain regions, and represent a crucial part of the human language network (Catani et al., 2005; Catani & Mesulam, 2008; Turken & Dronkers, 2011). This underlines the close relation of language and praxis processes (e.g., Kertesz & Hooper, 1982; Goldenberg & Randerath, 2015; Weiss et al., 2016). The central role of white matter damage observed in the present study strengthens the assumption that a brain network is involved in praxis skills and it is in line with previous studies. For example, Kertesz and Ferro (1984) reported that smaller lesions that induce apraxia are predominantly found in the periventricular white matter. Also, a combined fMRI-DTI study in healthy subjects identified a fronto-temporo-parietal network to underlie the ability to pantomime (Vry et al., 2015).

The finding of a network underlying apraxia is convincing since the ability to pantomime object use is a complex task that requires a wide range of cognitive and motor abilities. Several cognitive models of praxis skills have been proposed in line with findings in apractic patients (e.g., Barbieri & de Renzi, 1988; Cubelli et al., 2000; Bartolo et al., 2003; Johnson-Frey, 2004; Frey, 2008; Jax et al., 2014; Goldenberg, 2017). Although there is no consensus on the cognitive model underlying pantomime (Goldenberg, 2017), the different models generally assume paths along multiple cognitive processes that lie between phonological or visual analysis of the input stimuli (e.g., the word ‘tooth brush’ or a picture of a tooth brush) and the motor response. For example, a classical cognitive model of apraxia assumes that gestures such as pantomime are conceptualised, converted into a motor programme, and then executed (e.g., Liepmann, 1908; Barbieri & De Renzi, 1988; Jax et al., 2014). More recent models (e.g. Buxbaum & Randerath, 2018) show even higher complexity, and include, among others, an action working memory and the processing of spatial relationships, which can take place in different coding frames (Buxbaum, 2001). Given such complex cognitive models, a disruption of different cognitive functions could induce deficits in pantomime. These deficits may also show different characteristics with differently affected cognitive subcomponents. Accordingly, errors in apraxia can qualitatively differ and dissociate (e.g., Buxbaum, 2001; Halsband et al., 2001; Goldenberg, 2011; Manuel et al., 2013; Finkel et al., 2018). Thus, previous studies so far might have mapped different aspects of pathological behaviour functions at once. Indeed, it has been shown that the neural correlates of different apractic error types can dissociate (Manuel et al., 2013; Finkel et al., 2018). It is thinkable that such subcomponent analyses provided deeper insights into the network underlying apraxia by isolating cognitive subfunctions. If these subfunctions are located in smaller modules, previous univariate analyses might have been more powerful in identifying these smaller modules of the network. Still, it is not known if multivariate analysis of these subcomponents might also reveal more complex neural correlates than previous univariate studies did.

The findings in the present study are not only able to reconcile several discrepancies within the lesion-behaviour mapping literature in apraxia, but also discrepancies between lesion-behaviour mapping studies and fMRI studies. As in lesion studies, fMRI experiments that investigated pantomime also found several different left hemisphere regions to be involved (for reviews see Johnson-Frey, 2004; Lewis, 2006; Niessen et al., 2014; see also Lausberg et al., 2015; Vry et al., 2015; Martin et al., 2016; Chen et al., 2017). Parietal regions, including the intraparietal sulcus, inferior parietal lobe, and/or superior parietal lobe, were found activated in nearly all studies during pantomime (Niessen et al., 2014). Beyond, activation in middle and inferior frontal gyrus, inferior, middle and superior temporal lobe, inferior occipital gyrus, precentral gyrus, and insula were reported in some of these studies (Lewis, 2006; Niessen et al., 2014; Lausberg et al., 2015; Martin et al. 2016). The present finding suggesting a complex network to underlie higher-order motor control thus is in line with these observations derived from healthy subjects.

### Limitations of the present study

Albeit the present results resolve some inconsistencies in the field, open questions also remain. Our analyses found critical voxels in hippocampus, amygdala, and the temporal pole. There are several possible explanations. First, signal in hippocampus and amygdala might actually originate from adjacent white matter tracts, i.e. the inferior occipital and the uncinate fascicles. In both white matter atlases (Bürgel et al., 2006; Thiebaut de Schotten et al., 2011) used in our study, the uncinate fascicle even shows considerable overlap with the hippocampus and amygdala, as mapped by the AAL atlas. A second explanation is that these findings represent an artefact of MLBM, as patients with damage to the hippocampus consistently show larger lesions that also affect temporal regions (cf. Goldenberg & Randerath, 2015). Like mass-univariate analyses, multivariate analyses appear to be prone to errors if damage systematically co-occurs between two or more regions (Sperber et al., in press), i.e. if statistical independence of damage to voxels/brain regions is violated. In other words, the dependence of voxels in regards to damage, i.e. collateral damage between voxels, is a biasing factor in SVR-LSM like in univariate analyses. Similarly, it has been shown for SCCAN that the method is slightly superior to mass-univariate analyses with respect to this bias, but still not perfect (Pustina et al., 2018). Note that there are two types of dependencies between voxels: i) dependence of collateral damage, and ii) functional dependence, as for example in brain networks where multiple brain regions contribute to a cognitive function (see Mah et al., 2014; Xu et al., 2018; Sperber et al., in press). Previous simulation studies found a superiority of multivariate methods only in regard to functional dependence between voxels (see Sperber et al., in press for further discussion).

Furthermore, the data used in the present study were collected over many years, and lesions could not have been delineated or normalised with algorithms that are up to date these days (cf. de Haan & Karnath, 2018). The use of the data base, however, comes with a major advantage: the data set is very large, and larger than most other data sets in the field of lesion behaviour mapping. This advantage was very crucial to us: it has been postulated that multivariate modelling of lesion data requires large data sets (Mah et a., 2014), and in a recent study we have shown that SVR-LSM is optimally based on at least 100-120 patients. Similarly, comparisons between sample sizes in a simulation study suggested that performance of SCCAN in smaller sample sizes is also not optimal; and although not explicitly tested, accuracy of results markedly increased at least up to 100 subjects (Pustina et al., 2018).

### Perspective and Conclusions

Our multivariate analyses strongly support the assumption of a fronto-temporo-parietal network underlying apraxia of pantomime. In a next step, one should further elaborate how this network is involved in other deficits of higher-order motor skills, such as apraxia of imitation or apraxia of real tool use. The present study as well as a first study that investigated apraxia of imitation using a multivariate region-pair approach (Achilles et al., 2017) suggest that multivariate lesion-behaviour mapping contribute to improving our understanding of these symptoms. It may deepen our knowledge on how brain regions in the network and their cognitive functions work together, and could allow us to take the next steps in understanding human motor cognition both from a neuroscientist’s and a clinician’s perspective. Nevertheless, the use of multivariate methods in lesion-behaviour mapping only emerged recently (Smith et al., 2013), and they still need further elaboration and optimisation.

## Acknowledgements

This work was supported by the Deutsche Forschungsgemeinschaft (KA1258/15-1), by the Friedrich Naumann Foundation (stipend to C.S.), and the Luxembourg National Research Fund (FNR/11601161 to D.W.). There are no conflicts of interest to report.

